# *Drosophila melanogaster* miPEP8 regulates cell size through its interaction with ref(2)P/p62

**DOI:** 10.1101/2025.07.17.665293

**Authors:** Carine Duboé, Clémence Guillon, Jessica Boutet, Nathanael Jariais, Carole Pichereaux, Karima Chaoui, Emmanuelle Näser, Jean-Philippe Combier, Christine Brun, Yvan Martineau, Odile Burlet-Schiltz, Serge Plaza, Bertrand Fabre

## Abstract

MiPEPs are microproteins encoded by primary transcripts of microRNAs (pri-miRNAs). Initially identified in plants, we recently characterized a miPEP in *Drosophila melanogaster*, named miPEP8, which is involved in the regulation of wing size. However, mechanisms at play are unknown. In the present study, we take advantage of the drosophila cell line Schneider 2 (S2) to further investigate miPEP8 function at the molecular level. Overexpressing miPEP8 in S2 cells induced a reduction of cell size as well as an increase of the proportion of cells in the G1 phase of the cell cycle and an increase of the autophagic flux. A proteomics analysis revealed that miPEP8 overexpression in S2 cells induces the upregulation of several proteins including the autophagosome cargo protein ref(2)P (the orthologue of the human p62/Sequestosome 1 protein). The interactome of miPEP8 was generated and revealed interactions between this miPEP8 and the mTORC1/autophagy pathway. Bioinformatics analysis identified a short linear motif (SLiM) on miPEP8 sequence. Mutation of this SLiM prevented the interaction between ref(2)P/p62 and miPEP8. Mutation of the SLiM also reverted the smaller cell size phenotype observed when overexpressing miPEP8 in S2 cells. Finally, the cell size phenotype was reversed when cells were treated with RNA interference targeting ref(2)P/p62, suggesting that this protein plays a role in regulating the cell size in Drosophila.

## Introduction

The discovery of the pervasive translation of short open reading frames (sORFs) into thousands of microproteins has revolutionized our view of the proteome (1, 2). These small (typically less than 100 amino acids) unannotated proteins are produced from a variety of RNAs in all organisms from the kingdom of life (3–7). Microproteins have appeared as important regulators of key cellular processes (8). Although thousands of microproteins have been identified across many model organisms, we still lack a comprehensive characterization for most of their functions (8). One of the organisms that pioneered the study of microproteins is *Drosophila melanogaster*, notably through the discovery of the function of the Pgc and Pri/Tal peptides (9–12). Although the proteome of *Drosophila melanogaster* is well characterized (13–19), the discovery of its microproteome is still in its infancy (3, 11, 20–22). Recent studies in this model organism show that many microproteins can be produced from the translation of sORFs in alternative reading frames within canonical ORFs (3, 20), or that some of them are encoded by microRNA (*miR*) genes (3, 23, 24). These microproteins produced from *miR* genes are called miPEPs (miRNA-encoded peptides) and were first discovered in plants (25). Since their discovery, it was shown that miPEPs are present in a variety of plants but also in animals (26), although they seem to carry different functions in these two kingdoms (26, 27). In plants, miPEPs were shown to increase the expression level of their corresponding pri-miRNAs (25). In animals, however, miPEPs play different molecular roles, likely through interaction with different proteins, albeit in some cases affecting similar pathways than their corresponding miRNAs (26, 28).

Our group recently identified a microprotein encoded by *miR-8* of *Drosophila melanogaster*, named miPEP8 (24). *MiR-8* was shown to regulate body size in Drosophila, including wings (29). This is mediated through the binding of the *miR-8* miRNA to the mRNA coding for the U-shaped (USH) protein, an inhibitor of the phosphoinositide-3 kinase (PI3K) activity (29). Our previously published data shows that miPEP8 is also involved in the regulation of wing size in flies, albeit with reduced phenotype compared to *miR-8* mutants (in which both the miRNA stem loop and the miPEP8 ORF are deleted) (24, 29). Interestingly, both the overexpression and mutation of miPEP8 induce a reduction in wing size in flies (24), suggesting that the correct dosage is necessary within cells. Overexpression of miPEP8 results in a lower allelic frequency ratio compared to controls, indicating potential issues with fecundity and viability, and suggesting the involvement of miPEP8 in fitness, a marker/phenotype used in population genetics for a long time that has emerged as a sensitive detector of gene function (30, 31). The molecular mechanisms involved in the reduction of wing size upon miPEP8 overexpression or deletion are still not elucidated (24). In the present study, we use the drosophila cell culture line Schneider 2 (S2) to further investigate miPEP8 function at the molecular level. Overexpressing miPEP8 in S2 cells induced a reduction of cell size, in agreement with the phenotypes observed in flies, as well as an increase of the proportion of cells in the G1 phase of the cell cycle and an increase of the autophagic flux. A mass spectrometry-based proteomics approach revealed that overexpressing miPEP8 in S2 cells induces the upregulation of several proteins including the autophagosome cargo protein ref(2)P (the orthologue of the human p62/Sequestosome 1 protein). In order to gain some insight on the mode of action of miPEP8, its interactome was generated and revealed interactions between this microprotein and the mTORC1 (mechanistic target of rapamycin complex 1)/autophagy pathway. Bioinformatics analysis identified a short linear motif (SLiM) on miPEP8 sequence. Mutation of this SLiM prevented the interaction between miPEP8 and several proteins, including ref(2)P/p62. Mutation of the SLiM also reverted the smaller cell size observed upon overexpression of miPEP8 in S2 cells. Finally, the small cell size phenotype was also abolished when cells were treated with RNA interference targeting ref(2)P, suggesting that this protein is involved in the function of miPEP8 in regulating S2 cell size in drosophila. Altogether, our data points towards a function of miPEP8 in the regulation of cell size of drosophila through its interaction with the autophagosome cargo protein ref(2)P.

## Experimental Procedures

### S2 Cell Culture

S2 cells were cultured as described previously (24).

### Confocal Microscopy

For imaging experiments, S2 cells were co-transfected using an actin-GAL4 driver with UAS-EGFP, UAS-EGFP-miPEP8, UAS-EGFP-miPEP8mt (mutation of the MOD_Plk1/MOD_Plk4 SLiM identified on miPEP8, the “REKSIL” amino acids were modified to “AAAAAA”) or UAS-miPEP8. S2 cells were transfected with effectene (Qiagen) as described in (24). Forty-eight hours after transfection, the cells were prepared for microscopy analysis as previously described (3). Briefly, cells were transferred on cover slides, fixed using 4% paraformaldehyde (Sigma-Aldrich) in phosphate buffer saline (PBS) during 20 min at 20°C, washed twice with PBS and mounted with ProLong™ Diamond Antifade Mountant (Invitrogen). Slides were analyzed on a SP8 Leica confocal microscope. Cell size was manually measured in pixel using the image J software (32). For lysotracker experiments, viable cells were incubated on the cover slides for 20 min in complete Schneider′s Insect Medium with 50 nM LysoTracker™ (ThermoFisher Scientific). Cells were washed with PBS before fixation and further fixed and mounted as described above. In the LysoTracker™ treated cells, the signal intensity was measured with Integrated Density “RawIntDen” (sum of the values of the pixels in the selection) and divided by the cell size.

### Cell cycle analysis

S2 cells transfected with UAS-EGFP or UAS-EGFP-miPEP8 were treated for 24h either with 1µM Hydroxyurea + 10µM Aphidicolin, 1.7µM 20 Hydroxyecdysone or 2.7µM Colcemid to synchronize cells respectively in S, G2 and M phase. The G1 phase was obtained by subtracting the areas of the other phases of the cell cycle.

For flow cytometry experiments, S2 cells were collected in 15mL tube. They were fixed in 1% formaldehyde on ice for 30 minutes. Then the cells were permeabilized in PBT (phosphate buffer saline + 0.1% Tween-20) at room temperature for 5 minutes. Nucleus were stained by adding 1µg/mL DAPI to the cell suspension for 30 minutes in the dark. Cell were then sorted using a LSRFortessa X-20 flow cytometer (BD Biosciences) and analysis was performed using the Flowjo^TM^ V10 software (BD Biosciences).

### Proteome analysis

S2 cells were transfected with UAS-EGFP or UAS-EGFP-miPEP8 before cell sorting. Cells were fixed in 1% formaldehyde on ice for 30 minutes, permeabilized in PBT at room temperature for 5 minutes and nucleus stained with 1µg/mL DAPI for 30 minutes. Then cells were sorted using a FACSAria Fusion flow cytometer (BD Biosciences). After cell sorting, cells were lysed in an SDS-based buffer (Tris 50mM pH 7.5, 5% SDS) and BCA assay (Thermo Scientific™) was performed. 10 µg of proteins were loaded on SDS-PAGE and in-gel digestion was performed as previously described (33). The resulting peptides (50ng) were analyzed by nanoLC-MS/MS using an UltiMate 3000 RS nanoLC system (ThermoFisher Scientific) coupled to a TIMS-TOF SCP mass spectrometer (Bruker). Peptides were separated on a C18 Aurora column (25cm x 75µm ID, IonOpticks) using a gradient ramping from 2% to 20% of B in 30 min, then to 37% of B in 3min and to 85% of B in 2min (solvent A: 0.1% formic acid in H2O; solvent B: 0.1% FA in acetonitrile), with a flow rate of 150nL/min. MS acquisition was performed in DIA-PASEF mode on the precursor mass range [400-1000] m/z and ion mobility 1/K0 [0.64-1.37]. The acquisition scheme was composed of 8 consecutive TIMS ramps using an accumulation time of 100ms, with 3 MS/MS acquisition windows of 25 Th for each of them. The resulting cycle time was 0.96 seconds. The collision energy was ramped linearly as a function of the ion mobility from 59 eV at 1/K0=1.6Vs cm−2 to 20 eV at 1/K0=0.6Vs cm−2. Resulting .d files were analyzed using DIANN version 2.0 (34) with default parameters except that the number of maximum variable modifications was set to 1, precursor charge range were set from 2 to 4, precursor m/z range was set from 400 to 1000, fragment ion m/z range was set from 100 to 1700 and cross-run normalization was set to global. A database containing proteins from Uniprot (July 2020, 42,675 sequences), miPEP8 and EGFP sequences and common contaminants was used. Proteins with a Log_2_ fold change < −1 or > 1 and a p-value from a welch t-test < 0.05 were considered differentially regulated.

The western-blot analysis validating the increased level of ref(2)P upon miPEP8 overexpression in S2 cells was performed using anti-ref(2)P (Abcam), anti-GAPDH (35) and anti-GFP antibodies (Cell Signaling Technology).

### Interactome analysis

S2 cells were transfected with UAS-EGFP, UAS-EGFP-miPEP8 or UAS-EGFP-miPEP8mt and harvested 48h after transfection. Cells were lysed in a buffer containing 50mM Tris pH 7.5, 50mM NaCl, 5% glycerol, 0.1% Igepal and protease inhibitor cocktail (Sigma Aldrich, Burlington, MA, USA) and immunoprecipitation was performed using 40 µl of slurry of ChromoTek GFP-Trap® Agarose beads (ChromoTek GmbH, Planegg-Martinsried, Germany) O/N at 4°C. Beads were washed four times in lysis buffer and boiled in 50 μl of 2X Laemmli for 10 minutes at 100°C. Proteins were then alkylated using 60 mM of chloroacetamide and in gel digestion was performed as previously described (33). The resulting peptides were injected either on a nanoRS UHPLC system coupled to a Thermofisher LTQ Orbitrap Velos or a Thermofisher Q Exactive plus as previously described (36). Raw data were analysed using MaxQuant version 1.6.14.0 (37) using default parameters and the same database as for proteome analysis. To determine the miPEP8 interactome, a ratio was for each protein calculated between the minimal intensity value measured among the EGFP-miPEP8 immunoprecipitation (IP) replicates and the maximal intensity value measured among the EGFP IP replicates as described in (38). The minimum ratio threshold to define a protein as a miPEP8 interacting partner was defined as previously described (39). Briefly, this threshold corresponds to Q3 + 1.5 × IQ, Q3 being the third quartile and IQ being the interquartile, both based on the distribution of all the Log_2_ ratio (EGFP-miPEP8 IP / EGFP IP) measured for each protein quantified in the experiment. In addition, a minimum ratio (EGFP-miPEP8 IP / EGFP IP) of 2 based on spectral counts was required to consider a protein as a miPEP8 partner. Finally, in the case of proteins present only in miPEP8 IPs (immunoprecipitations) samples, an intensity superior to the Q1 (first quartile, based on the distribution of all the measured intensity) had to be observed in at least one of the miPEP8 IP replicates to consider the protein as a miPEP8 partner.

In the case of the comparison of the interactome of EGFP-miPEP8mt and EGFP-miPEP8, proteins were considered differentially associated if they were identified as interactor of miPEP8 (see above) and the Log_2_ ratio of the average intensities of EGFP-miPEP8mt on the average intensities of EGFP-miPEP8 was < −1 or > 1 and the p-value from a welch t-test < 0.05. Gene Ontology terms enrichment and protein networks were generated using String v12.0 (40).

### Data Availability

All the mass spectrometry data have been deposited with the MassIVE repository with the dataset identifier: MSV000098269.

### miPEP8 interactors validation by co-immunoprecipitation

Plasmids coding for HA-tagged versions of interactors identified by MS as described above (and the GAPDH as a negative control) were obtained from the Berkeley Drosophila Genome Project (BDGP) (41). S2 cells were co-transfected with UAS-EGFP, UAS-EGFP-miPEP8 or UAS-EGFP-miPEP8mt and the plasmid for each protein interactor (and the GAPDH as a negative control) and a western-blot analysis was performed using anti-HA (Sigma-Aldrich) and anti-GFP antibodies (Cell Signaling Technology).

### Split-Luciferase analysis

S2 cells were cultured in 24 wells plate for 18h before transfection. Then they were co-transfected with plasmids containing an actin driver and miPEP8-N-ter Luc (N-terminal fragment of the luciferase) and the C-ter Luc (C-terminal fragment of the luciferase) fused in N-ter of the different candidates. For the competition condition, a plasmid containing an actin-GAL4 driver and the candidate’s sequences were also transfected. S2 cells were transfected with effectene (Qiagen) according to the manufacturer’s instructions. Forty-eight hours after transfection, cells were transferred in a white 96 wells plate to measure luminescence by adding 5µL of 10mM D-luciferin. Kinetics of luminescence emission were monitored using the Victor Nivo (Perkin Elmer) plate reader.

### Plasmids, dsRNAs and qPCR

The dsRNAs were generated with respectively pUFO Rag C/D-HA (DGRC number UFO02654), pUFO Ref(2)P-FLAG-HA (generated by subcloning Ref(2)P sequence, DGRC number FMO06145) and Luciferase ICE T7 control vector (Promega) according to (41) using the following primers as described in the Supplementary table 1. S2 cells were harvested 48h post-transfection. RNAs were extracted using the RNeasy kit (Qiagen) and reverse transcription were performed using the M-MLV Reverse Transcriptase (Promega) following manufacturer’s instructions. qPCR analysis was performed using the LightCycler 480 SYBR Green I Master (Roche Life Science) on a LightCycler 480 Instument II (Roche Life Science) and analyzed with the LightCyler 480 Software (Roche Life Science), using the 2-ΔΔCt method (42). A list of the primers used can be found in Supplementary table 1.

### Short linear motif identification and miPEP8 interactor predictions

Short linear motifs (SLiMs) have then been detected in the miPEP8 sequence using the ELM prediction tool of the ELM database (43), considering only the true positive instances detected in D. melanogaster. mimicINT, a workflow initially developed for microbe-host protein interactions inference (44) has been used to infer protein-protein interactions between miPEP8 and the Drosophila canonical proteins. Adapted to the sPEP-canonical protein in Drosophila, it then consists in (i) the detection of short linear motifs, i.e. SLiMs, extracted from the ELM database (43) and globular domains predicted using the existing InterPro signatures (45) in the sPEP sequence; (ii) the collection of domains on the Drosophila canonical proteins; (iii) the interaction inferences between sPEP and canonical proteins, according to interaction templates (see (44) for further details). Notably, only true positive instances of SLiMs detected in *D. melanogaster* are considered for interaction predictions (http://elm.eu.org/instances/?q=&instance_logic=&taxon=Drosophila+melanogaster).

### Detection of protein domains on miPEP8 sequence

InterProScan (v5.52-87.0)(46) was used to look for protein domains on miPEP8 sequence.

## Results and discussion

### Overexpression of miPEP8 in S2 cells decreases cell size, accumulates cell in G1 phase and changes the expression of a set of proteins

In order to gain some insights into the molecular function of miPEP8, we used Schneider 2 (S2) cells as it is one of the most used cell lines for studies in drosophila. MiPEP8 was not detected in this cell line (24), thus we decided to work with overexpression of this microprotein in S2 cells. Upon overexpression of an EGFP-miPEP8 (enhanced green fluorescent protein-miPEP8) fused protein, we observed a decrease of S2 cell size (Figure 1A-B). The median size of cells overexpressing EGFP-miPEP8 was 26.8 % smaller than the median size of cells overexpressing EGFP (p = 8.3E-10) (Figure 1B). This diminution of cell size is in agreement with previous data showing that overexpressing miPEP8 in drosophila wings induces a decrease of their size (24). To avoid any artifacts linked to the fusion of miPEP8 (71 amino acid long) to an EGFP protein, we monitored the effect of the transfection of a plasmid expressing a version of miPEP8 not fused to any tag on cell size. Overexpression of miPEP8 promoted a decrease in S2 cell size (p = 3.8E-06) similar to the one induced by the overexpression of EGFP-miPEP8 (Supplementary Figure 1), showing that miPEP8 retains its activity even fused with a bigger protein (EGFP). Flow cytometry analysis showed that overexpression of miPEP8-EGFP accumulates cells in the G1 phase of the cell cycle compared to cells overexpressing EGFP (p = 2.9E-02) or non-transfected cells (p = 3.4E-03) (Supplementary Figure 2).

**Figure 1:**
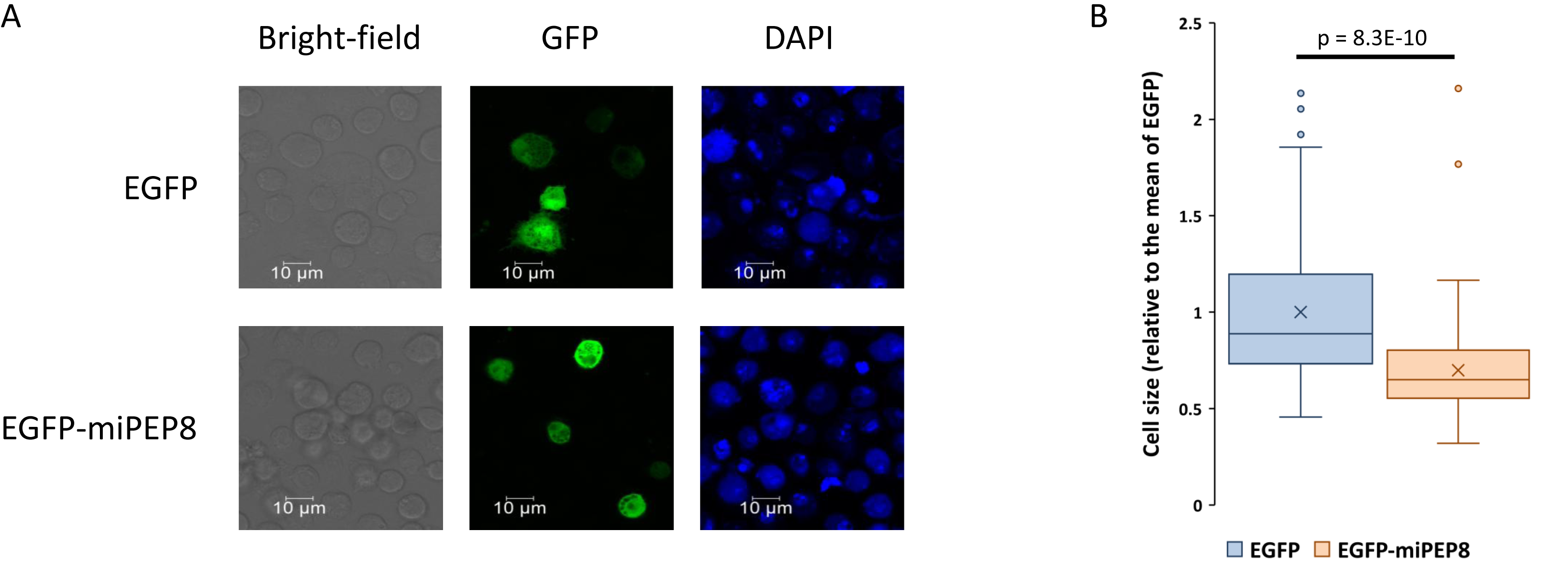
Overexpression of miPEP8 decreases S2 cell size. A. S2 cells were transfected with a plasmid encoding either the EGFP or EGFP fused to miPEP8 (EGFP-miPEP8). Cells were stained with DAPI and observed by confocal microscopy. B. The size of the S2 cells expressing either EGFP or EGFP-miPEP8 was measured and normalized to the mean of the measured EGFP cell size. The p-value was calculated using a Welch’s t-test (n = 100 and 108 for EGFP and EGFP-miPEP8, respectively).

The expression of miPEP8 in S2 cells induces a phenotype, albeit discrete, on cell size and cell cycle (Figure 1A-B, Supplementary Figure 1-2). Thus, we decided to monitor how the overexpression of miPEP8 alters the proteome of S2 cells. S2 cells were transfected with either EGFP as a control or EGFP-miPEP8, cells expressing either protein were sorted using fluorescence-activated cell sorting (FACS), lysed, proteins were digested with trypsin and analyzed in data independent acquisition (DIA) on a TIMS-TOF SCP mass spectrometer (Bruker) (Figure 2A). 5,451 proteins were quantified, 29 proteins being differentially regulated between EGFP and EGFP-miPEP8 expressing cells (2 more abundant in EGFP and 27 more abundant in EGFP-miPEP8 expressing cells) (Figure 2B and Supplementary table 2). Among the proteins up-regulated (Figure 2C) we could find an enrichment of proteins involved in cellular response to stress (p = 2.5E-02) and more particularly cellular response to heat stress (p = 6E-04) (Supplementary Figure 3). This set of proteins includes small heat shock proteins (Hsp22, Hsp23, Hsp26 and Hsp27), the expression of which is developmentally regulated and involved in a wide range of function (17, 47). In addition, the Atg18b (autophagy related gene 18b) and ref(2)P (refractory to sigma P) proteins, both involved in autophagy (48), were also found increased following overexpression of EGFP-miPEP8 (Figure 2B-C). This increase in protein expression was confirmed for ref(2)P by western-blot (Figure 2D). Reanalyzing previously published RNA-seq data (24), we could observe an increase of the mRNA levels of the small heat shock proteins (Hsp22, Hsp23, Hsp26 and Hsp27) upon overexpression of miPEP8 but not for ref(2)P, suggesting a post-transcriptional mechanism for the regulation of this protein (Supplementary Figure 4). In summary, this proteomics analysis points towards a possible role of miPEP8 in the regulation of cellular stress.

**Figure 2:**
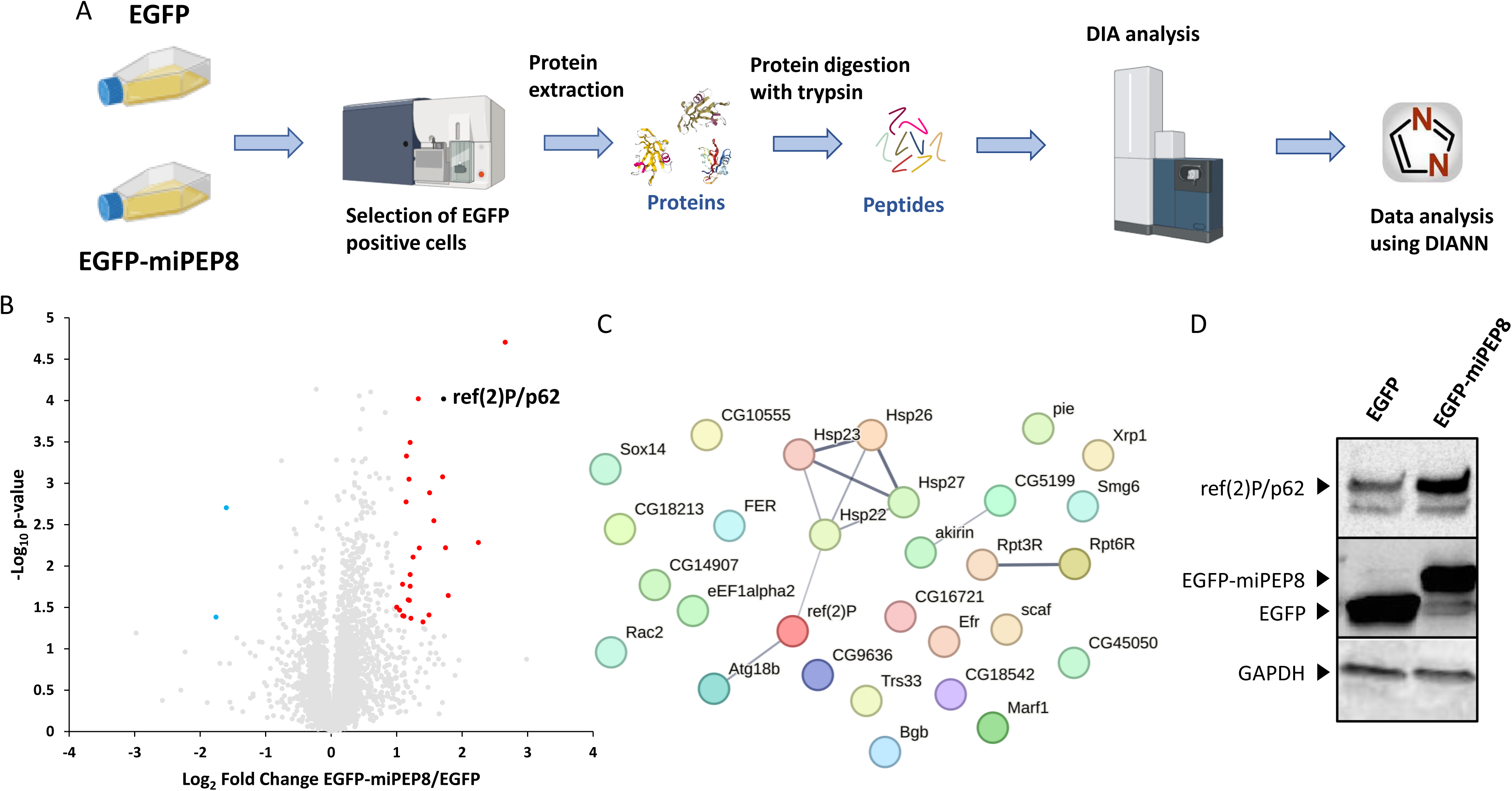
Data independent analysis proteomics workflow to identify proteins differentially regulated in S2 cells upon overexpression of miPEP8. A. S2 cells were transfected with EGFP or EGFP-miPEP8 ( n = 3 each) and the cells expressing the fluorescent protein were selected by fluorescence-activated cell sorting. Proteins were extracted and digested with trypsin and resulting peptides were run in data-independent acquisition mode on a TIMS-TOF SCP mass spectrometer (Bruker). The resulting .d files were analyzed with DIANN. B. Volcano plot representing the log_2_ ratio (EGFP-miPEP8/EGFP) for each protein quantified and the corresponding p-value obtained from a Welch’s t-test. The blue, red, and gray dots represent the proteins more abundant in the cells expressing EGFP, more abundant in the cells expressing EGFP-miPEP8, or not differentially expressed between the two conditions, respectively. The protein ref(2)P/p62 is displayed in black on the volcano plot. C. STRING network of the proteins more abundant in the EGFP-miPEP8 condition. The line thickness between two proteins indicates the strength of data support. D. Western blot analysis for ref(2)P/p62, GFP and GAPDH were performed on the lysates of S2 cells expressing EGFP or EGFP-miPEP8.

### Defining miPEP8 interactome in S2 links this microprotein with the intracellular signaling and autophagy pathways

Given that the overexpression of miPEP8 impacts cell size and induces the expression of proteins involved in cellular stress response, the interactome of this microprotein was generated to identify its potential protein partners (Figure 3A). Either the EGFP or the EGFP-miPEP8 fusion was overexpressed in S2 cells and, after cell lysis, an affinity purification was performed using GFP-Trap agarose beads (Figure 3A). Following protein digestion, peptides were analyzed by LC-MS/MS (Figure 3A) and stringent criteria were used to consider a protein as a miPEP8 interacting protein (see methods section). Despite these stringent criteria, 211 proteins were found interacting with miPEP8 (Figure 3B and Supplementary table 3) suggesting that miPEP8 might be involved in several cellular pathways (Figure 3B-C). Among the most represented pathways, Signaling, Cell cycle, Cell division, Regulation of MAPK cascade, mTOR signaling, Autophagy and Apoptosis were identified (Figure 3B-C). All these pathways were shown to affect cell size in some capacity (49–51). Several of these pathways, namely mTOR, apoptosis and autophagy are linked to the cellular stress response, as observed in the proteome of miPEP8 overexpressing cells (Supplementary Figure 3).

**Figure 3:**
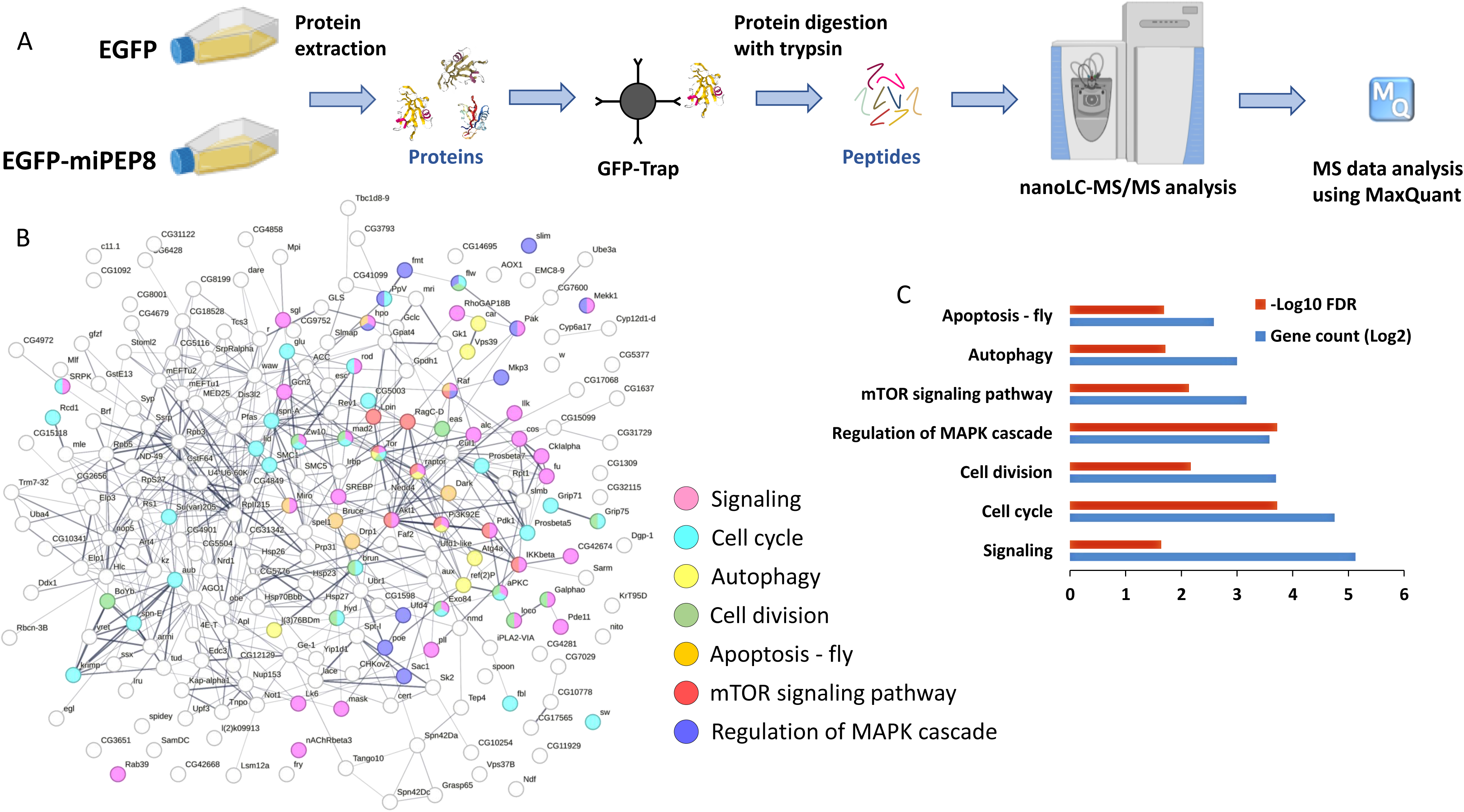
The interactome of miPEP8 reveals its interaction with a wide range of cellular pathways. A. S2 cells were transfected with EGFP or EGFP-miPEP8 (n = 3 for EGFP-miPEP8 and n = 4 for EGFP) and proteins were extracted and subjected to a GFP-Trap agarose beads (Chromotek) precipitation. After elution, proteins were concentrated in one band on SDS-PAGE and digested with trypsin and resulting peptides were run in data-dependent acquisition mode on a Q Exactive plus mass spectrometer (Thermofisher). The resulting raw files were analyzed with MaxQuant. B. STRING network of the proteins found as interacting with EGFP-miPEP8. The proteins belonging to the pathways found enriched by STRING were displayed with different colors. The line thickness between two proteins indicates the strength of data support. C. GO term analysis using STRING for the proteins interacting with miPEP8. The number of proteins identified in each pathway (gene count, log2 transformed) and the corresponding p-value (−log10 transformed) are represented on the graphs.

Co-immunoprecipitation (IP) experiments were used to validate the interaction between miPEP8 and protein candidates such as Gcn2 (general control nonderepressible 2), Atg4a (autophagy related gene 4a), PI3K92E (Phosphatidylinositol 3-kinase 92E) and RagC-D (Ras-related GTP binding C/D) (Figure 4A). S2 cells were co-transfected with plasmids encoding these candidates fused with a HA tag and either the EGFP or EGFP-miPEP8 proteins. Immunoprecipitation using GFP-Trap agarose beads (Chromotek) was performed followed by western blot analysis. For all the candidates tested a signal was observed in the EGFP-miPEP8 IP but not in the EGFP IP, confirming that all these proteins indeed interact with miPEP8 (Figure 4A). GAPDH-HA (glyceraldehyde-3-phosphate dehydrogenase) was used as negative control to ensure that miPEP8 was not binding its partners in a non-specific manner. Contrary to the other candidates tested, we could not detect any interaction between miPEP8 and GAPDH, strengthening our interactome data (Figure 4A). As a complementary approach, we used split luciferase complementation assays to validate some of the interaction observed by MS (Figure 4B). In this assay, two interacting candidates (here miPEP8 and one of its interacting partners) are both fused to a different fragment of the firefly luciferase (typically referred as N-terminal and C-terminal fragments)(52). If the interacting candidates bind, the fragments of the firefly luciferase are brought into proximity, thus enabling the reconstitution of the enzyme and detection of a luminescent signal can be observed upon addition of its substrate luciferin (52). One of the advantages of protein-fragment complementation assays over co-immunoprecipitation experiments is that the distance required for the fragment of the reporter protein (here the firefly luciferase) needs to be short (inferior to 100 Å) meaning that the candidate interacting proteins are in close proximity and thus this assay reports direct or very close interactions (52). Here, miPEP8 and the candidates were fused to the N-terminal fragment or the C-terminal fragment of the firefly luciferase, respectively (Figure 4B). Each construction encoding the candidates fused to the C-terminal fragment of the firefly luciferase was co-transfected in S2 cells with the construction encoding miPEP8 fused to the N-terminal fragment of the reporter enzyme. In order to check that both fragments of the firefly luciferase did not interact on their own in our experimental conditions, plasmids encoding each fragment not fused to any protein were co-transfected (Figure 4B). Upon lysis of the cells and addition of luciferin, a small luminescence signal was measured (Figure 4B). On the contrary, as a positive control of interaction, the proteins Eyeless (Ey) and Antennapedia (Antp) were fused to N-terminal fragment and the C-terminal fragment of the firefly luciferase, respectively (Figure 4B). These two proteins were previously shown to interact (53). After lysis of the cells and addition of luciferin, a luminescence signal 14.5 time higher than that of the negative control was measured (p = 1.7E-03) (Figure 4B) validating our split luciferase approach. Looking at the candidates, luminescence signals were observed for all the proteins tested with a minimal signal 4.6 times higher than the negative control in the case of PI3K92E (p = 7.3E-04) (Figure 4B). In order to check the specificity of the observed interaction, we decided to implement a competition experiment (Figure 4B). Plasmids encoding the candidates tagged with a HA motif (Comp) were co-transfected in S2 cells with the construction encoding the candidates fused to the C-terminal fragment of the firefly luciferase and miPEP8 fused to the N-terminal fragment of the reporter enzyme. The rationale behind the addition of the HA tagged candidates is that if the HA tagged proteins compete with the candidates fused to the C-terminal fragment of the firefly luciferase for the interaction with miPEP8, a decrease in the luminescence signal should be observed, attesting for the specificity of the interaction between miPEP8 and its partners. A significant reduction in the luminescence signal in the presence of the competitor (Comp) was observed for all the tested candidates, with the less striking decrease observed for PI3K92E (2.8-fold less measured signal upon addition of the competitor, p = 7E-04) (Figure 4B). Overall, these experiments show that miPEP8 interacts with several proteins involved in intracellular signaling pathways and autophagy (Figure 3B-C and Figure 4A-B).

**Figure 4:**
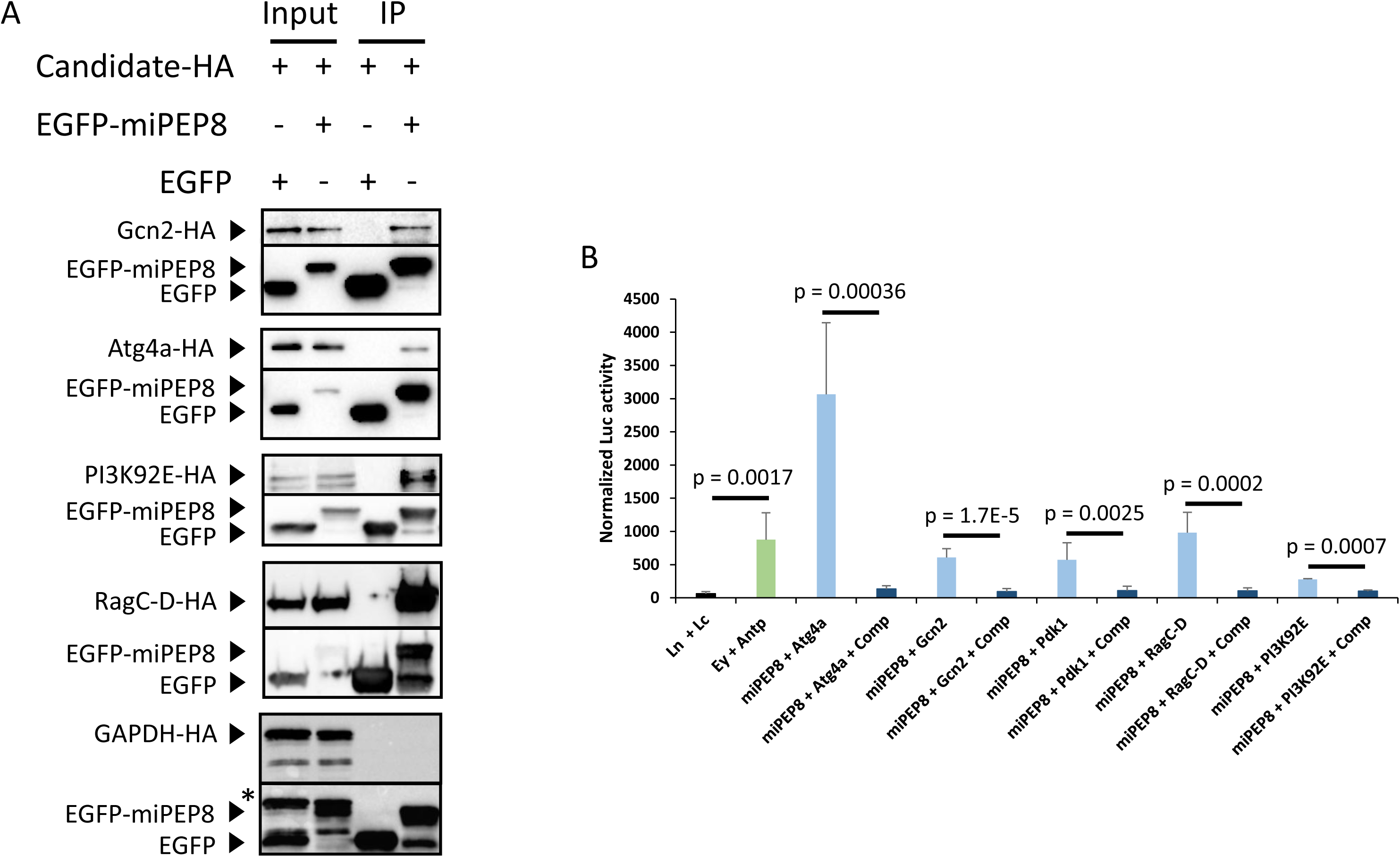
Validation of the interaction between miPEP8 and proteins involved in the autophagy and intracellular signaling pathways. A. Co-immunoprecipitation experiments were performed after lysis of S2 cells transfected with plasmids encoding miPEP8 interacting candidates fused with a HA tag and plasmids encoding either EGFP or EGFP-miPEP8 (n = 3 for each candidate). Western blot analysis for HA and GFP were performed. B. S2 cells were transfected with plasmids expressing miPEP8 fused with the N-terminal fragment of the luciferase and each protein candidate fused with the C-terminal fragment of the luciferase. In the case of addition of a competitor (Comp), S2 cells were also transfected with a plasmid encoding the corresponding candidate fused with a HA tag (and not fused to the C-terminal fragment of the luciferase). The N-terminal (Ln) and C-terminal (Lc) of the luciferase not fused to any proteins were used as negative control. The proteins eyeless (Ey) and Antenapedia (Antp) were used as positive control of interacting proteins. The p-value was calculated using a Welch’s t-test (n = 7 for each candidate except PI3K92E for which n = 3).

### miPEP8 bears a short linear motif responsible for its interaction with ref(2)P/p62

To better understand how miPEP8 functions at the molecular level, we looked for the presence of potential protein domains on its sequence using InterProScan (46). Aside from a predicted disordered region spanning from the amino acid 36 to 71, no protein domain could be identified in the sequence of miPEP8. The presence of Short Linear Motifs (SLiMs) was then investigated. SLiMs are short amino acid sequences generally found in disordered regions and involved in protein-protein interactions (54). Using the ELM database SLiM prediction, several potential short motifs were identified on the sequence of miPEP8 (Supplementary Figure 5A). Based on the SLiMs predicted on the miPEP8 sequence, interactions between miPEP8 and canonical proteins from *Drosophila melanogaster* were inferred using the mimicINT workflow (44). The predicted and experimental interactors (based on our interactome data) were compared and a list of 19 proteins emerged as found in both approaches (Supplementary Figure 5B). All these 19 proteins were protein kinases (Supplementary Figure 5B-C) and were predicted to bind the MOD_Plk1/ MOD_Plk4 SLiM spanning from the residues 21 to 26 on miPEP8 sequence (Figure 5A and Supplementary Figure 5A). Structural prediction of miPEP8 using AlphaFold (55) showed a central position of this SLiM within this microprotein (Figure 5B). Despite the fact that the MOD_Plk1/ MOD_Plk4 SLiM is supposed to be a predicted phosphorylation site on the serine 24 of miPEP8, we could not observe any phosphorylation on this amino acid in our MS data, only the non-phosphorylated serine 24 was observed (Supplementary Figure 6). We then decided to monitor the effect of the mutation of this SLiM motif (replaced by an alanine stretch, Figure 5A) on the interactions between miPEP8 and its partners. An affinity purification followed by MS analysis was performed using the GFP-Trap agarose beads (Chromotek) as described above but S2 cells were transfected with either EGFP, EGFP-miPEP8 or EGFP-miPEP8mt (mutated SLiM MOD_Plk1/ MOD_Plk4). The mutation of the MOD_Plk1/ MOD_Plk4 SLiM changed the interaction of 26 proteins with miPEP8 (Figure 5C and Supplementary table 4). Among the proteins differentially interacting, one stood out, ref(2)P, the drosophila homolog of the human protein p62/sequestosome-1, involved in autophagy (48) and found upregulated in the proteome of S2 cells upon overexpression of miPEP8 in S2 cells (Figure 2A-C). Upon mutation of the MOD_Plk1/ MOD_Plk4 SLiM, the interaction between miPEP8 and ref(2)P/p62 decreased by a 2.8 fold (p = 0.0022) (Figure 5C and Supplementary table 4). This change in interaction was confirmed by western-blot analysis of a co-immunoprecipitation experiment in which S2 cells were co-transfected with ref(2)P-HA and either EGFP, EGFP-miPEP8 or EGFP-miPEP8mt (Figure 5D). In parallel, we also co-transfected another interactant of miPEP8, Gcn2 (Figure 3B, Figure 4A-B and Supplementary table 3), that was not affected by the mutation of the MOD_Plk1/ MOD_Plk4 SLiM in our MS data (Figure 5C and Supplementary table 4), and either EGFP, EGFP-miPEP8 or EGFP-miPEP8mt followed by co-immunoprecipitation and western-blot analysis (Figure 5D). As observed in our MS interactome data (Figure 5C and Supplementary table 4), the mutation on the SLiM did not impair the interaction between miPEP8 and Gcn2, confirming that the MOD_Plk1/ MOD_Plk4 SLiM is involved in the interaction with only a subset of proteins (Figure 5D).

**Figure 5:**
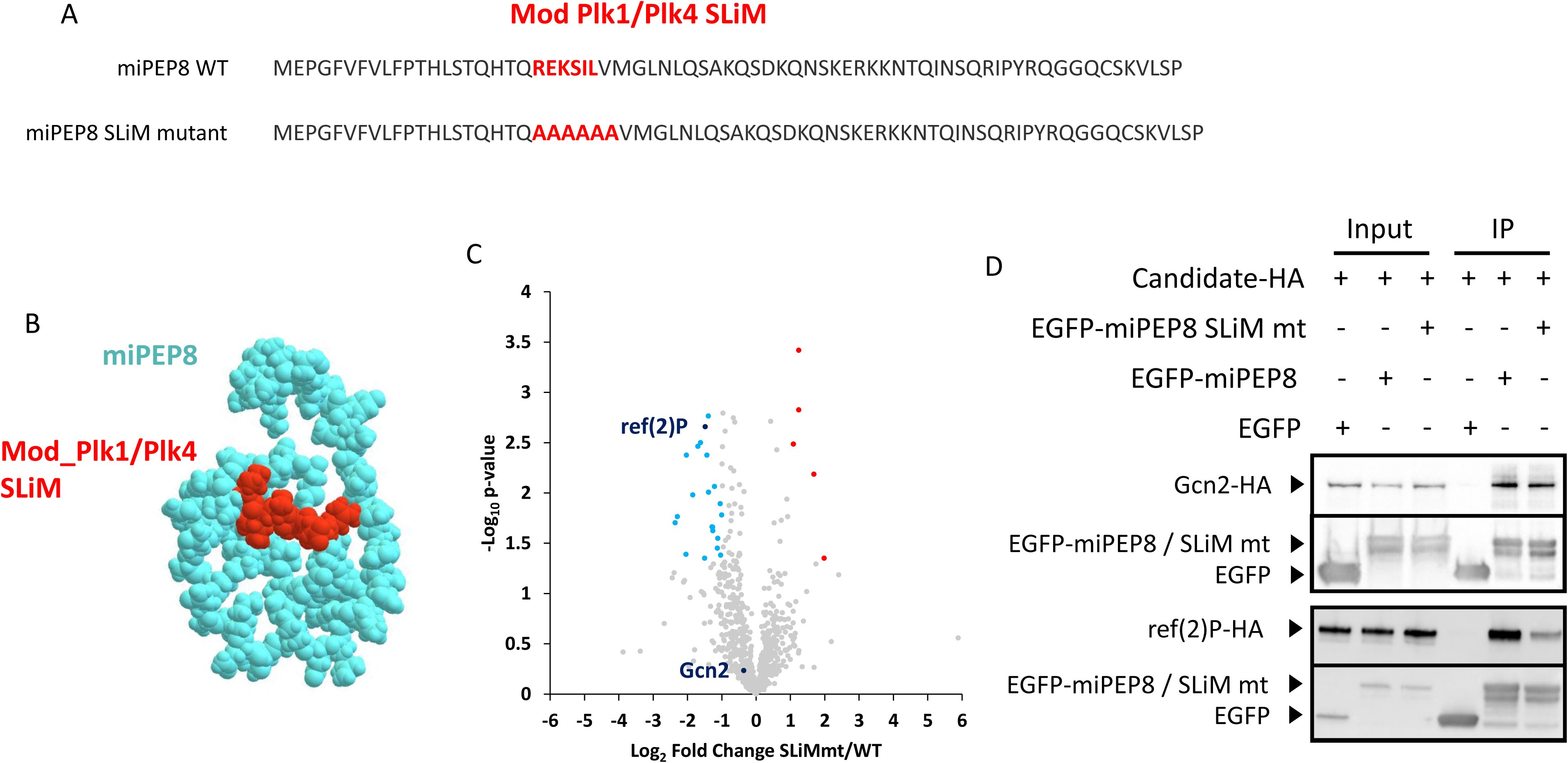
A MOD_Plk1/ MOD_Plk4 SLiM on miPEP8 is involved in its interaction with the protein ref(2)P/p62. A. Amino acid sequence of miPEP8 and its mutant. The sequences corresponding to the MOD_Plk1/ MOD_Plk4 SLiM and the resulting sequence in the mutant (miPEP8mt) are shown in red. B. Predicted structure of miPEP8 generated using AlphaFold. C. Volcano plot representing the log_2_ ratio (EGFP-miPEP8mt/EGFP-miPEP8) for each protein quantified in the EGFP-miPEP8mt or EGFP-miPEP8 immunoprecipitations (IPs) and the corresponding p-value obtained from a Welch’s t-test (n = 4). The blue, red, and gray dots represent the proteins more abundant in the EGFP-miPEP8 IP, more abundant in EGFP-miPEP8mt IP, or not differentially associated between the two conditions, respectively. The proteins ref(2)P/p62 and Gcn2 are displayed in dark blue on the volcano plot. D. Co-immunoprecipitation experiments were performed after lysis of S2 cells transfected with plasmids encoding ref(2)P-HA or Gcn2-HA and plasmids encoding either EGFP, EGFP-miPEP8 or EGFP-miPEP8mt (n = 3). Western blot analysis for HA and GFP were performed.

### Mutation of the MOD_Plk1/Plk4 SLiM on miPEP8 or knocking down ref(2)P/p62 expression reverse the decreased of S2 cell size observed upon overexpression of miPEP8

The effect of the mutation of the MOD_Plk1/ MOD_Plk4 SLiM on miPEP8 on cell size was then investigated to see if this motif is involved in the reduction of S2 cell size observed upon overexpression of miPEP8 (Figure 1A-B). S2 cells were transfected with either EGFP, EGFP-miPEP8 or EGFP-miPEP8mt (Figure 6A). As expected, a decreased size of S2 cells expressing EGFP-miPEP8 was observed compared to cells expressing EGFP (p = 3E-03) (Figure 6A). However, cells transfected with EGFP-miPEP8mt had a similar size to EGFP transfected cells (p = 9.5E-01) and were bigger than cells expressing EGFP-miPEP8 (p = 6E-03) (Figure 6A). This result points towards a role of the MOD_Plk1/ MOD_Plk4 SLiM on miPEP8 on the decreased cell size observed upon overexpression of this microprotein. We then decided to monitor if proteins that are interacting with miPEP8 through this motif were also involved in the phenotype smaller cells. The protein ref(2)P was first investigated as it seems to interact with miPEP8 via its MOD_Plk1/ MOD_Plk4 SLiM (Figure 5C-D) and was found upregulated in cells overexpressing miPEP8 (Figure 2A-C). Double stranded RNA (dsRNA) targeting either the RNA of luciferase protein (dsLuc, negative control) or the RNA of ref(2)P (dsref(2)P) through RNA interference were added to the culture medium of S2 cells for 72 hours. Cells were then transfected with either EGFP or EGFP-miPEP8 and the cell size of EGFP positive cells was measured. We controlled that an important decrease of the mRNA level of ref(2)P was observed upon treatment with its dsRNA (Supplementary Figure 7A). Despite the presence of dsLuc, overexpression of EGFP-miPEP8 still induced a decrease in cell size compared to EGFP (p = 0.001), showing that application of a control dsRNA does not affect the miPEP8-induced phenotype (Figure 6B). The dsref(2)P also induced a significant reduction of EGFP transfected cell size compared to cells treated with dsLuc (p = 9E-03) (Figure 6B). Surprisingly, despite the fact that both overexpression of miPEP8 and silencing of ref(2)P result in a reduction of the size of S2 cells, the combination of the expression of the microprotein and down-regulation of ref(2)P reversed the observed phenotype back to control level (EGFP dsLuc, p = 9.1E-01) (Figure 6B). Such reversal could not be observed when using dsRNA targeting another protein, RagC-D (Supplementary Figure 7B), another protein interacting with miPEP8 (Figure 3B, Figure 4A-B and Supplementary table 3), despite a good efficiency of RagC-D mRNA silencing (Supplementary Figure 7C). Overall, these results indicate that miPEP8 interacts with ref(2)P and controls S2 cell size through this interaction.

**Figure 6:**
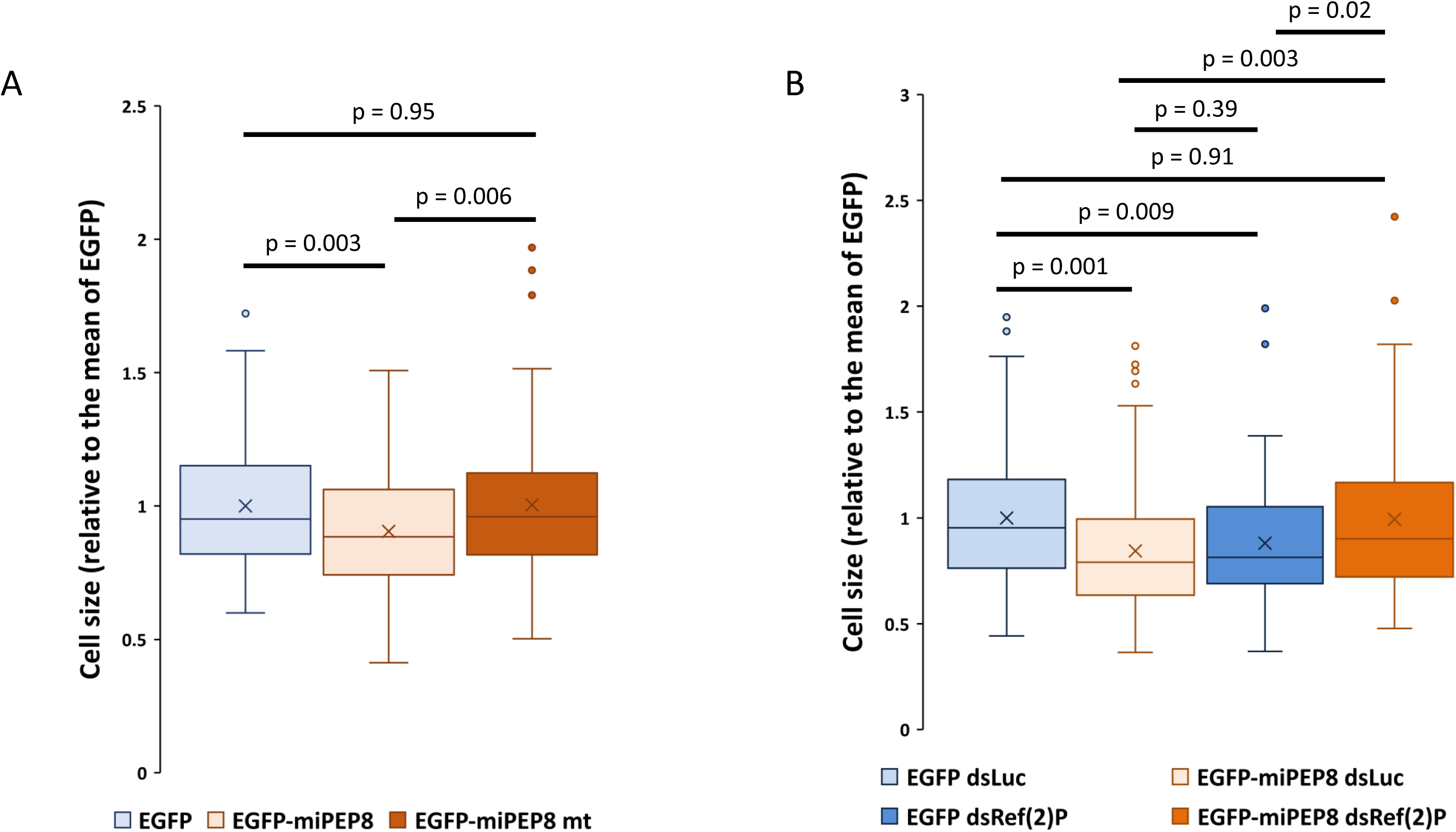
Mutation of the MOD_Plk1/ MOD_Plk4 SLiM on miPEP8 or knocking down ref(2)P expression reverse the reduction of S2 cell size observed upon overexpression of miPEP8. A. The size of the S2 cells expressing either EGFP, EGFP-miPEP8 or EGFP-miPEP8mt was measured and normalized to the mean of the measured EGFP cell size. The p-values were calculated using a Welch’s t-test (n = 89, 89 and 84 for EGFP, EGFP-miPEP8 and EGFP-miPEP8mt, respectively). B. The size of the S2 cells expressing either EGFP or EGFP-miPEP8 and treated with either Luciferase (dsLuc) or ref(2)P (dsref(2)P) dsRNAs (RNA interference) was measured and normalized to the mean of the measured EGFP dsLuc cell size. The p-values were calculated using a Welch’s t-test (n = 83, 89, 90 and 91 for EGFP dsLuc, EGFP-miPEP8 dsLuc, EGFP dsref(2)P and EGFP-miPEP8 dsref(2)P, respectively).

As ref(2)P is involved in autophagy, we decided to monitor the effect of an overexpression of miPEP8 on the autophagic flux. We used the LysoTracker™ to assess the presence of lysosomes as previously described (21). A significant increase in the intensity of the signal could be noticed between cells expressing EGFP-miPEP8 versus cells expressing EGFP (Supplementary Figure 8). We then tested whether this increase of autophagic flux depends on the MOD_Plk1/ MOD_Plk4 SLiM. Surprisingly, the increase of the intensity of the LysoTracker™ signal was also observed in cells transfected with EGFP-miPEP8mt (Supplementary Figure 9). This result indicates that the modulation of the autophagic flux induced by the overexpression of miPEP8 is not directly linked to the concomitant increase of the protein level of ref(2)P (Figure 2B-D). MiPEP8 might rather regulate the autophagic flux in S2 cells in a mechanism independent of its MOD_Plk1/ MOD_Plk4 SLiM.

## Conclusion

As it has recently been shown, microproteins seem to have a wide range of functions in the development and physiology (fitness) of *Drosophila melanogaster* (21, 22). A previous study from our group identified a microprotein encoded by the *miR-8* gene than is involved in fly survival and wing size regulation (24). Here, we investigated how miPEP8 functions at the molecular level. Overexpression of this microprotein in S2 cells showed a robust decrease of cell size and alteration of the cell cycle (Figure 1 and Supplementary Figure 2). This is consistent with the phenotype of flies overexpressing miPEP8, which have smaller wings and a reduced allelic frequency ratio, indicating reduced survival capacity (24). Here, we show that the reduction of cell size seems to be mediated through the interaction between miPEP8 and ref(2)P/p62 via a SLiM located on miPEP8 as knocking down ref(2)P/p62 or mutating the SLiM on miPEP8 brought back the cells to their original sizes (Figure 6). In addition to the decrease of cell size, overexpression of miPEP8 in S2 cells also accumulated cells in the G1 phase of the cell cycle and increased the autophagic flux (Supplementary Figure 2 and 8). The latter is not mediated through the MOD_Plk1/ MOD_Plk4 SLiM and thus not depending on the interaction between miPEP8 and ref(2)P/p62 (Supplementary Figure 8). Among the SLiMs identified on miPEP8, one of them in the N-terminal part of the microprotein is a LIG_LIR_Gen_1 which is involved in the binding of proteins to the forming autophagosome (Supplementary Figure 5). In addition, the protein level of Atg18b was found upregulated upon miPEP8 overexpression and Atg4a was identified as a protein partner of miPEP8 (Figure 3B and Figure 4A-B). Both of these proteins are involved in autophagosome formation (56, 57). It would be interesting to see if these two proteins and the LIG_LIR_Gen_1 identified on miPEP8 are involved in the increased autophagic flux observed upon overexpression of miPEP8.

Besides its role in regulating cell size and autophagy, miPEP8 might have other function in drosophila. Indeed, despite the use of stringent criteria to identify proteins associated to miPEP8 in our interactome data, more than 200 proteins were found interacting with this microprotein (Figure 3B). Although we cannot exclude that some of these interactions might be due to the overexpression of miPEP8, it is quite surprising to observe many protein kinases associated to this small protein (Figure 3B and Supplementary table 3). Our data might point to a function of miPEP8 in the regulation of key signaling pathways such as the PI3K/Akt/mTORC1 or MAPK pathways (Figure 3B). Recent studies have suggested that many microprotein-encoding genes play a role in organismal fitness, as revealed by weak developmental defects and detectable phenotypes that only appear when organisms are challenged (21, 22). One possible explanation is that microproteins are modulators of cellular pathways. Our results support this hypothesis, as neither the gain-of-function nor the loss-of-function of miPEP8 induces strong developmental defects. Moreover, the interactors of miPEP8 identified are components of pathways involved in cell survival, fitness, and stress response (Figure 3B-C). All in all, the present study provides a first molecular insight into the function of miPEP8 in drosophila and precise its role in the regulation of cell size through an interaction with ref(2)P, as well as paving the way to future study of the involvement of this microprotein in other cellular pathways.

## Acknowledgments

This work has been supported by the Fondation ARC pour la recherche sur le cancer. This work was funded by the French ANR project AltFUNC (ANR-23-CE44-0028). The work was funded in part by grants from the Région Occitanie, European funds (Fonds Européens de Développement Régional, FEDER), Toulouse Métropole, and the French Ministry of Research with the Investissement d’Avenir Infrastructures Nationales en Biologie et Santé program (ProFI, Proteomics French Infrastructure project, ANR-10-INBS-08). This work was supported by the Institut National du Cancer (INCa 2021-073, Y.M.).

