## Supplementary legends for "*Drosophila melanogaster* miPEP8 regulates cell size through its interaction with ref(2)P/p62"

Supplementary figures legends

**Supplementary figure 1: Overexpression of miPEP8, fused or not with EGFP, decreases S2 cell size.** S2 cells were transfected with a plasmid encoding either the EGFP, EGFP fused to miPEP8 (EGFP-miPEP8) or two plasmids, one encoding EGFP and the other miPEP8 not fused to any tag (EGFP + miPEP8). Cells were stained with DAPI and observed by confocal microscopy. The size of the S2 cells expressing either EGFP, EGFP-miPEP8 or EGFP + miPEP8 was measured and normalized to the mean of the measured EGFP cell size. The p-value was calculated using a Welch's t-test (n = 95, 88 and 92 for EGFP, EGFP-miPEP8 and EGFP + miPEP8, respectively).

**Supplementary figure 2: Overexpression of EGFP-miPEP8 accumulates S2 cells in the G1 phase of the cell cycle.** S2 cells were transfected with a plasmid encoding either the EGFP or EGFP fused to miPEP8 (EGFP-miPEP8), then stained with DAPI and analyzed by flow cytometry. The p-value was calculated using a Welch's t-test (n = 6).

**Supplementary figure 3: STRING Reactome pathway analysis of the proteins upregulated upon overexpression of EGFP-miPEP8 in S2 cells.**

**Supplementary figure 4: Reanalysis of previously published RNAseq data showing the $Log_2$ fold change of of RNA expression of *ref(2)P*, *Hsp23*, *Hsp26* and *Hsp27* upon overexpression of miPEP8 in S2 cells.** The p-value was calculated using a Welch's t-test (n = 5).

**Supplementary figure 5: Identification of SLiMs motifs in miPEP8's amino acid sequence and prediction of protein interactors.** A. Short linear motifs (SLiMs) detected in miPEP8's sequence using the ELM prediction tool of the ELM database. The MOD_Plk_1/ MOD_Plk_4 SLiMs are highlighted in a red box B. String network of the proteins both predicted to interact with miPEP8 using mimicINT and identified in the miPEP8 interactome. The line thickness between two proteins indicates the strength of data support. C. String Molecular Function GO terms analysis of the proteins showed in (B).

**Supplementary figure 6: MS/MS spectra of the EKSILVMoxGLNLQSAK peptide from miPEP8 which includes the serine 24 not phosphorylated.**

**Supplementary figure 7: Efficiency of dsRNA targeting ref(2)P.** Real time quantitative PCR analysis of cells transfected with EGFP or EGFP-miPEP8 and treated with dsRNAs (RNA interference) targeting the luciferase protein (dsLuc, negative control) or ref(2)P (dsref(2)P).

**Supplementary figure 8: Knocking down RagC-D does not reverse the reduction of S2 cell size observed upon EGFP-miPEP8 overexpression.** A. The size of the S2 cells expressing either EGFP or EGFP-miPEP8 and treated with either Luciferase (dsLuc) or RagC-D (dsRagC-D) dsRNAs (RNA interference) was measured and normalized to the mean of the measured EGFP dsLuc cell size. The pvalues were calculated using a Welch's t-test (n = 93, 102, 95 and 95 for EGFP dsLuc, EGFP-miPEP8 dsLuc, EGFP dsRagC-D and EGFP-miPEP8 dsRagC-D, respectively). B. Real time quantitative PCR analysis of cells transfected with EGFP or EGFP-miPEP8 and treated with dsRNAs (RNA interference) targeting the luciferase protein (dsLuc, negative control) or RagC-D (dsRagC-D).

**Supplementary figure 9: Overexpression of EGFP-miPEP8 or EGFP-miPEP8mt in S2 cells increases the autophagic flux.** S2 cells were transfected with a plasmid encoding either the EGFP, EGFP-miPEP8 or EGFP-miPEP8mt and treated with LysoTracker™ to assess the presence of lysosomes. The LysoTracker™ intensities measured by confocal microscopy were normalized by the cell size for each condition. An ANOVA with Welch correction was performed and for groups comparison a Games-Howell post-hoc test was used (n = 458, 346 and 384 for EGFP, EGFP-miPEP8 and EGFP-miPEP8mt, respectively).

**Supplementary Table 1 : List of the primer sequences used for the reverse transcription-quantitative polymerase chain reaction experiments and list of the Ds RNAs used for RNA interference.**

**Supplementary Table 2 : Table containing the protein intensity measured by MS in S2 cells transfected with EGFP or EGFP-miPEP8.**

**Supplementary Table 3 : Table containing the proteins identified and quantified in the EGFP and EGFP-miPEP8 immunoprecipitations (IP) and proteins that are specifically enriched in the EGFP-miPEP8.**

**Supplementary Table 4 : Table containing the proteins identified and quantified in the EGFP, EGFP-miPEP8 and EGFP-miPEP8mt immunoprecipitations.**
