## Supplementary figures for "*Drosophila melanogaster* miPEP8 regulates cell size through its interaction with ref(2)P/p62"

Supplementary Figure 1

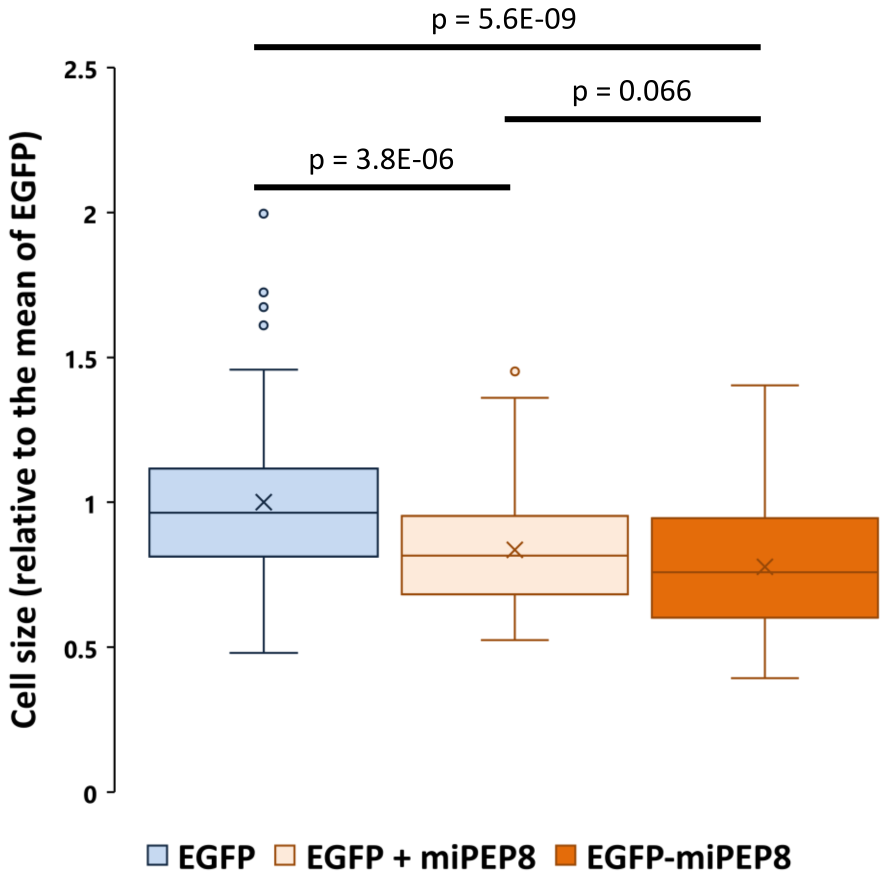

Supplementary Figure 2

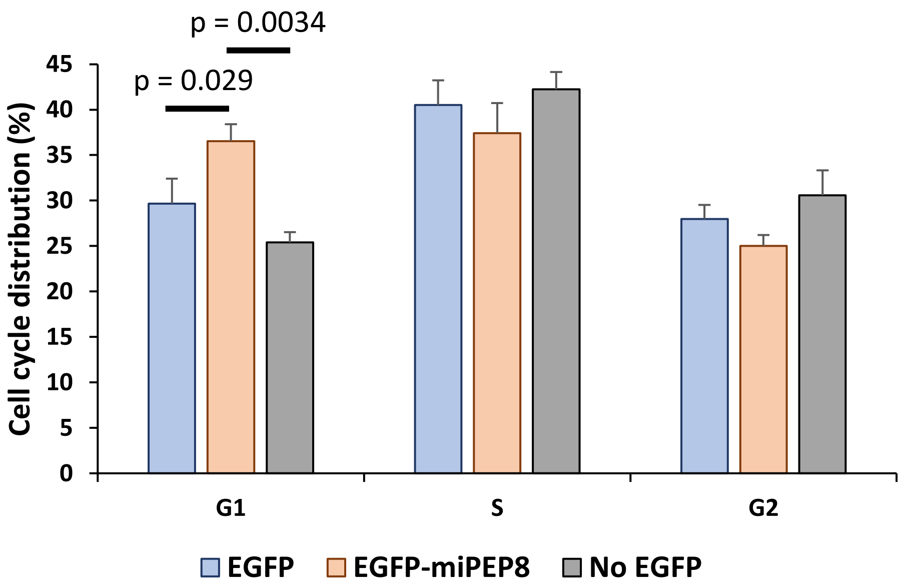

Supplementary Figure 3

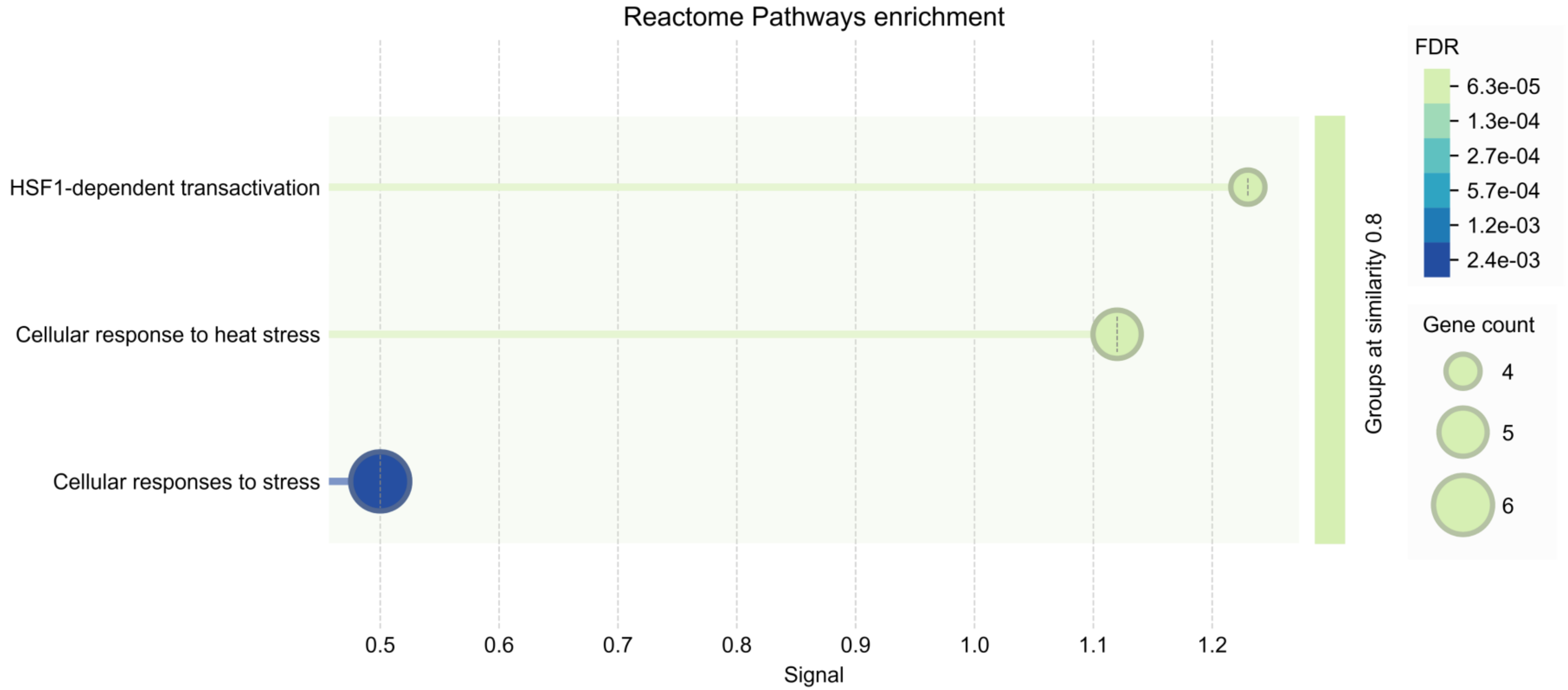

Supplementary Figure 4

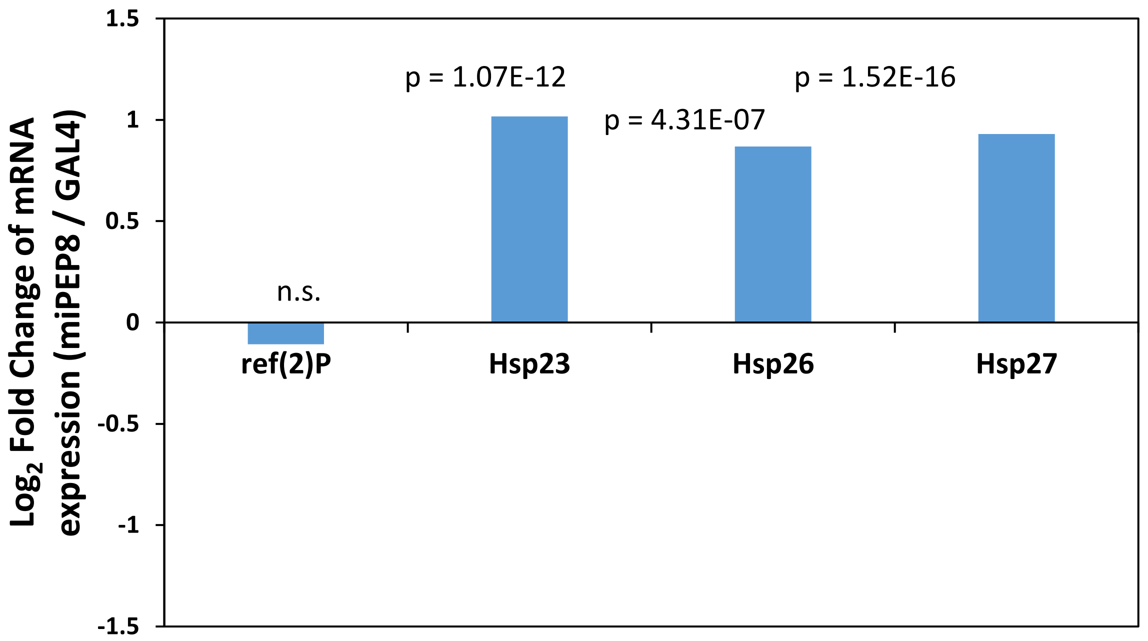

### Supplementary Figure 5

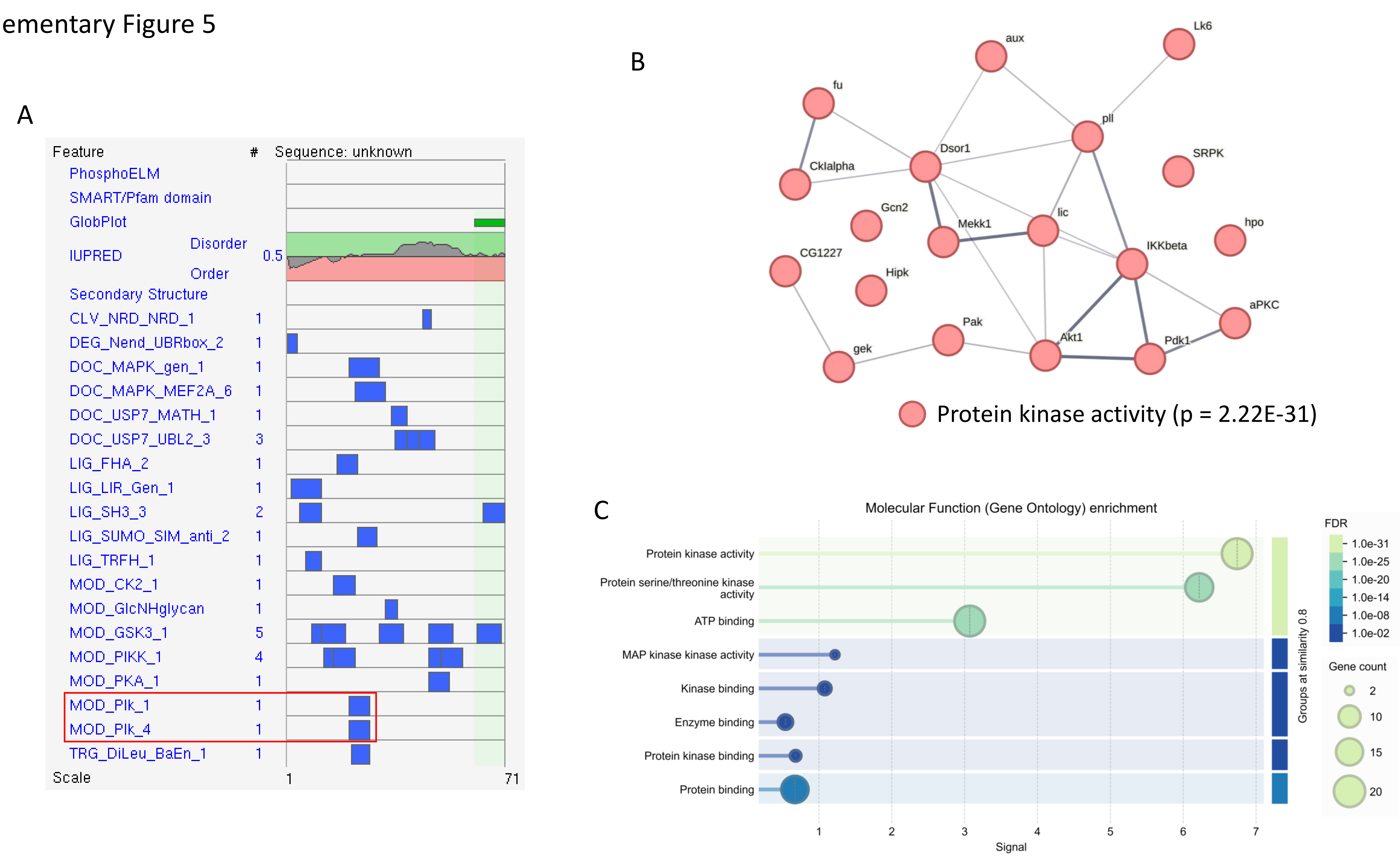

Supplementary Figure 6

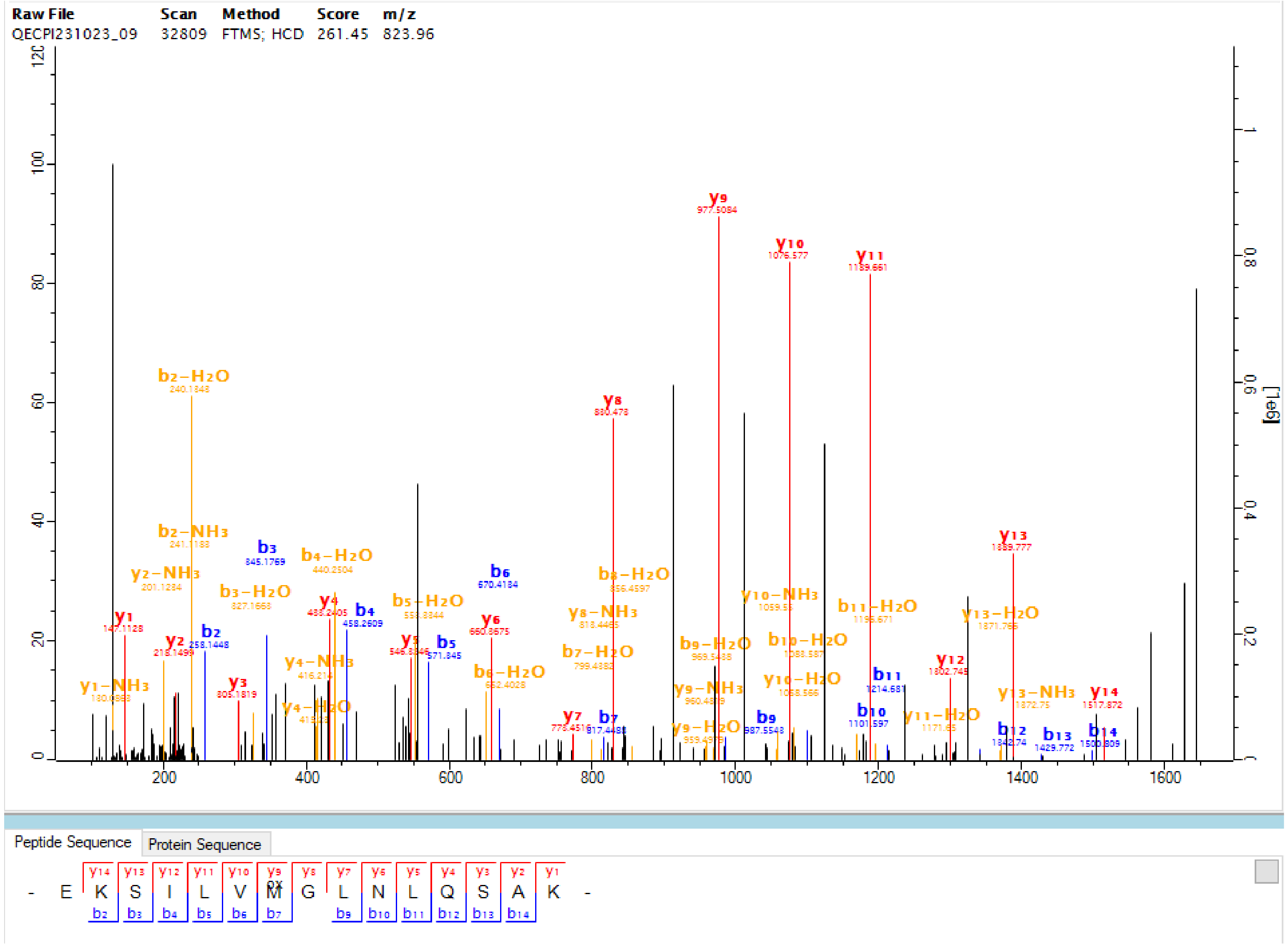

Supplementary Figure 7

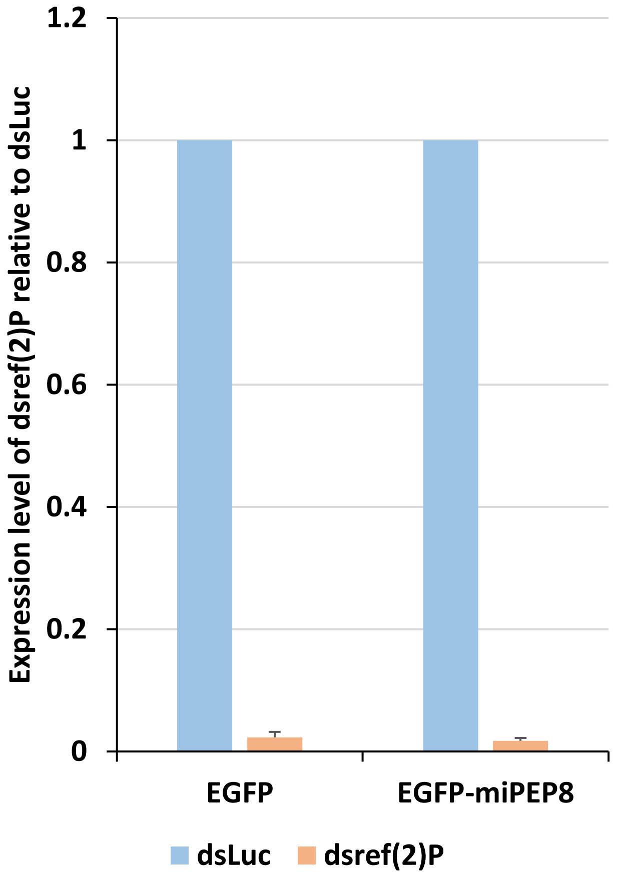

Supplementary Figure 8

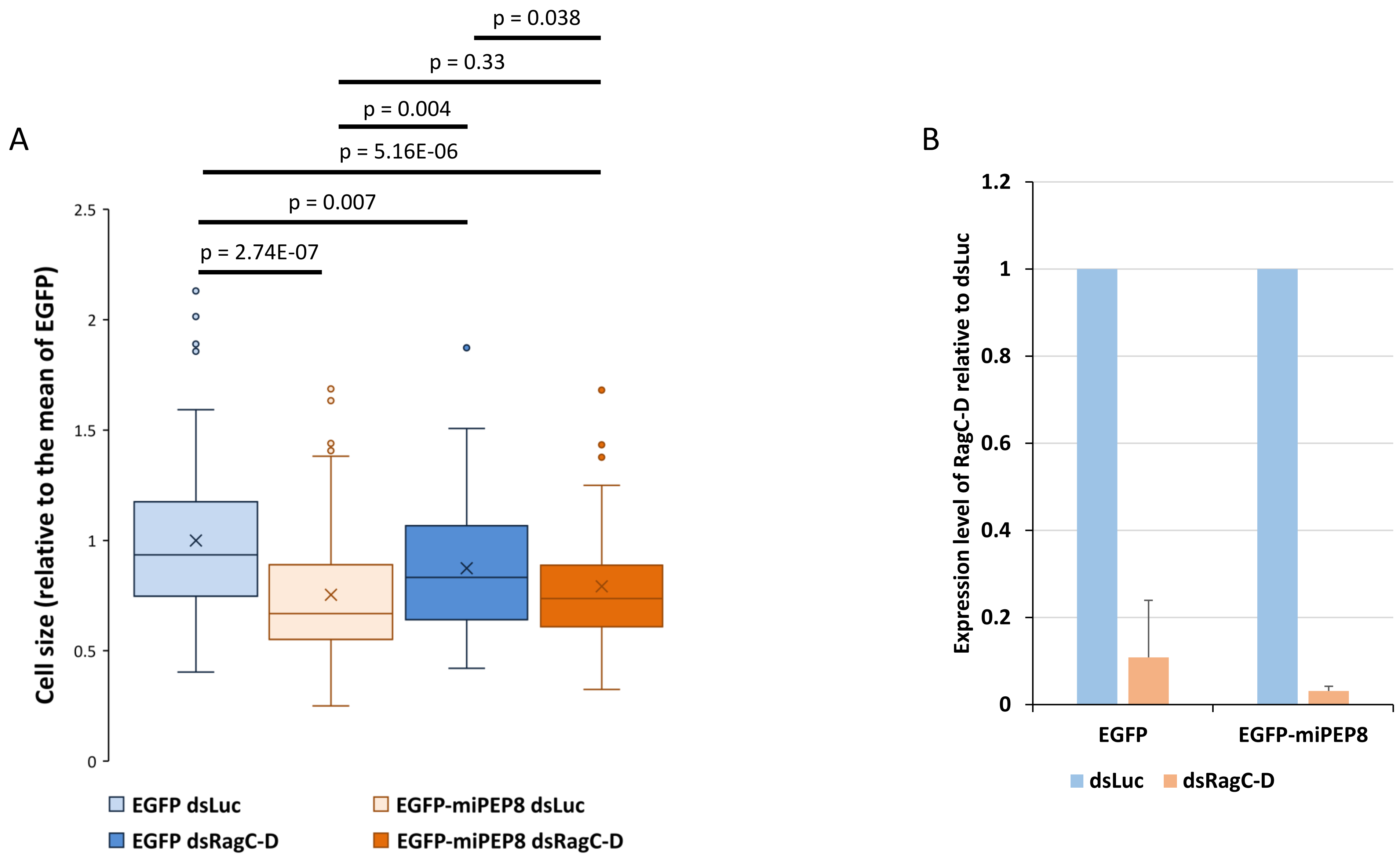

Supplementary Figure 9

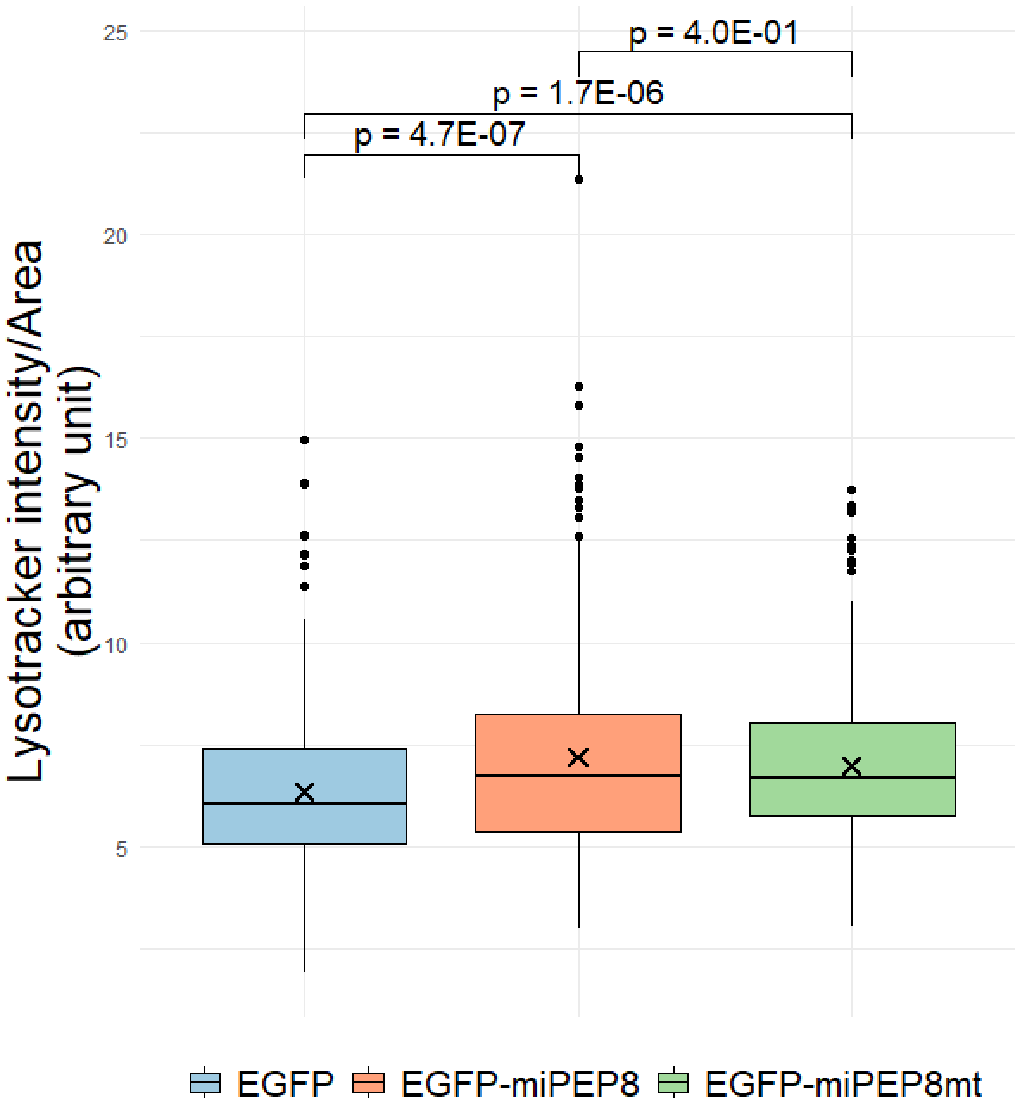
